# GrimACE: Automated, multimodal cage-side assessment of pain and well-being in mice

**DOI:** 10.1101/2025.03.07.642046

**Authors:** Oliver Sturman, Marcel Schmutz, Tom Lorimer, Runzhong Zhang, Mattia Privitera, Fabienne K Roessler, Justine Leonardi, Rebecca Waag, Alina-Mariuca Marinescu, Clara Bekemeier, Katharina Hohlbaum, Johannes Bohacek

## Abstract

Pain and welfare monitoring is essential for ethical animal testing, but current cage-side assessments are qualitative and subjective. Here we present the GrimACE, the first fully standardised and automated cage-side monitoring tool for mice, the most widely used animals in research. The GrimACE uses computer vision to provide automated mouse grimace scale (MGS) assessment together with pose estimation in a dark, safe environment. We validated the system by analysing pain after brain surgeries (craniotomies) with head implants under two analgesia regimes. Human-expert and automated MGS scores showed very high correlation (Pearson’s r=0.87). Both expert and automated scores revealed that a moderate increase in pain can be detected for up to 48 hours after surgeries, but that both a single dose of meloxicam (5mg/kg s.c.) or 3 doses of buprenorphine (0.1mg/kg) + meloxicam (5mg/kg s.c.) provide adequate and comparable pain management. Simultaneous pose estimation demonstrated that mice receiving buprenorphine + meloxicam showed increased movement 4h after surgery, indicative of hyperactivity, a well-known side-effect of opioid treatment. Significant weight loss was also detected in the buprenorphine + meloxicam treatment group compared to the meloxicam-only group. Additionally, detailed BehaviorFlow analysis and automated MGS scoring of control animals suggests that habituation to the GrimACE system is unnecessary, and that measurements can be repeated multiple times, ensuring standardised post-operative recovery monitoring.

## INTRODUCTION

The evaluation of pain and well-being in laboratory animals is an essential part of all ethical experimentation ^1,2^. Although mice are the most commonly used animal model in scientific studies due to their genetic similarities to humans and their utility in understanding various diseases and treatments ^3,4^, accurate assessment of pain and well-being in mice is often challenging. Inadequate pain management not only raises serious ethical concerns regarding the humane treatment of animals but also jeopardizes the validity and reproducibility of research findings ^5,6^. With good pain and welfare monitoring protocols in place it is possible to design appropriate analgesia regimes, detect problems before they become too severe and to define and work with humane endpoints.

Moderate to severe pain in laboratory mice is most often related to surgical interventions, which are a cornerstone of in-vivo animal research. The gold standard to assess post-surgical recovery is based on cageside assessment ^7^, where visual inspection of key behavioral parameters (e.g. posture, coat condition, movement patterns, wound licking) are assessed by a trained experimenter ^2,8–10^. These visual observations, which provide qualitative scores or counts for individual parameters, are well-suited for rapid assessment of post-surgical recovery by personnel trained in animal experimentation or by trained animal caretakers. However, cageside assessment is prone to bias, subjectivity and poor sensitivity to subtle alterations in well-being ^11^. It is also widely believed that prey animals may hide signs of pain, making it difficult for observers in close proximity to accurately assess subtle changes in well-being ^12^. In contrast, measures like telemetry for movement and heart rate, nest building behavior or burrowing behavior have been shown to be more sensitive indicators of post-surgical pain and recovery, which can reveal pain-related changes when standard cage-side assessment fails to reveal impairments ^13–17^. However, these tests initially require surgeries to implant transmitters or habituation and then prolonged observation periods in single-housing to analyze complex behaviors, rendering these tests impractical for routine use in laboratories that do not specialize in pain assessment. Over the last decade, the assessment of facial features to detect the affective component of pain has been popularized through the development of the mouse grimace scale (MGS) ^18,19^. This approach requires minimal habituation and only brief periods of surveillance using photo or video recordings. Subsequent manual scoring assesses whether signs of pain can be detected across five facial features (orbital tightening, nose bulge, cheek bulge, ear position and whisker change), and each feature is assigned a value from 0 to 2 (0=absent, 1=moderate, 2=severe). This scoring process is very labor intensive, requires highly trained experimenters ^18,20^, and remains subject to bias ^21^. Several groups have developed pipelines to automate (parts of) this process ^22–26^, but since experimental setups vary between labs, automated pipelines do not transfer well between labs and setups. Moreover, in many scenarios, the assessment of grimace scores with automated software is particularly challenging, for example, when head-implants or other interventions (fresh wound sites on the head with ointment from sterilization and local anesthetics) alter the images. This is particularly problematic, as craniotomies are the most commonly used surgical procedures in neuroscience research, and pain associated with craniotomies is notoriously difficult to detect using cage-side assessment ^27,28^. MGS scores are highly sensitive to pain after craniotomies, showing that pain typically peaks 4-6 hours after surgery, before resolving gradually over the course of 24-48 hours ^6,9,27,29^. These studies also suggest that non-steroidal anti-inflammatory drugs (NSAIDs) like meloxicam or carprofen provide adequate analgesia ^27,29^. Finally, going beyond classical pain assessment tools, deep behavioral profiling has recently emerged for pain detection, leveraging machine-learning to extract subtle behavioral motifs from video recordings of freely moving animals ^30–32^. These approaches were able to reveal subtle behavioral alterations in mouse pain models ^30^, resolve a broad range of effects following stress exposure^33^, and distinguish the impact of different pharmacological interventions ^34^. However, these tools are still difficult to implement in most labs, as they require substantial technical expertise ^35^.

Here, we present the GrimACE, a robust and portable cageside assessment system that minimizes stress levels (dark, safe environment), collects high-quality videos with full-body pose estimation for behavioral profiling, and acquires standardized high quality facial images with automated MGS scoring trained on data from expert human raters. On the pose estimation data we then apply deep behavioral phenotyping using our newly developed BehaviorFlow pipeline ^33^. We test the GrimACE device by assessing postoperative analgesia following craniotomies with different kinds of brain implants, comparing NSAID to opioid + NSAID analgesia. We show that from 10 min recordings, the GrimACE can automatically detect mild-to-moderate post-operative pain from facial features, when cageside assessment does not. Behavioral profiling readily reveals known side-effects of opioid treatment and together with the MGS tracks recovery over time.

## RESULTS

### The GrimACE

The GrimACE is a complete hardware and software solution for standardised MGS image acquisition, scoring, and keypoint-based pose estimation. The GrimACE consists of an aluminium frame with custom 3d printed (PLA) components to hold lights, a front and top camera, and an acrylic arena with similar dimensions to other MGS arenas ^24,29,36,37^ (Figure 1a,b). The arena has matte white walls and an infra-red (IR) permeable front and lid. The use of IR-permeable black acrylic on the front and lid of the box allows a clear view of the mouse from the front and above, creates a dark calming environment for the mouse, and ensures that the lighting of the room has minimal effect on the image/video quality. The arena also has small air holes to ensure that the mouse remains comfortable. The arena is mounted on an orange 3D-printed base that slides into a standardised position within the frame, ensuring repeatability. The matte white acrylic and orange PLA help to reduce reflections and provide good image contrast. Large matte white acrylic sheets attached to the aluminium frame help to reduce reflections from external light sources. The small acrylic box and 3d printed base have also been designed to fit together without glue or screws so that they can be dismantled and reassembled in seconds by hand, which makes them easy to clean. Once the mouse is in the arena and is in position, the user can trigger the system to begin recording video, acquiring and analysing images using the GrimACE app.

**Figure 1.**
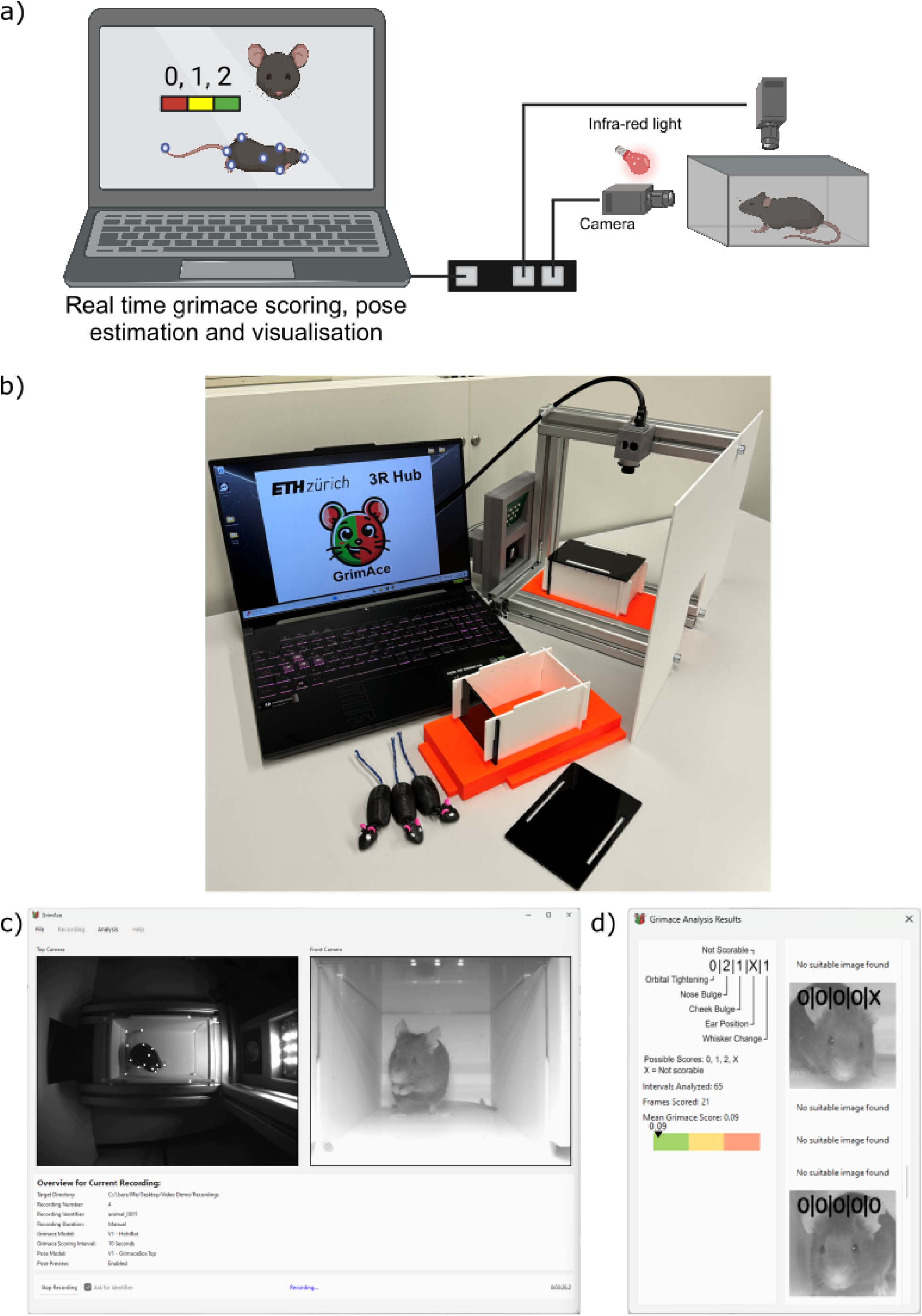
The GrimACE, a complete hardware and software solution for standardised MGS scoring and keypoint-based pose estimation. (a) Schematic of the GrimACE. (b) The GrimACE, with extra arena and 3d printed mice that can be used for demonstrative purposes (top acrylic shield has been removed to increase visibility for this photograph) (c) The GrimACE app during a recording, with the pose estimation video on the left and MGS video on the right. (d) The live MGS window, with scored images, the current mean score and detailed score breakdowns.

#### GrimACE app and machine learning algorithm overview

The GrimACE app provides a clean, simple, easy-to-use interface for running MGS scoring and pose estimation. The user sets the output file naming convention (e.g. an experiment name prefix), the desired test duration (e.g. 10 mins), and the MGS scoring interval (e.g. 10 seconds), and these settings persist from test to test. When the mouse is in position, the app is triggered manually, and displays several real-time quality checks throughout the test: front video, top video (including pose estimation) (Figure 1c), and cropped scored images with detailed score breakdowns (Figure 1d). At the end of each test, two videos (top and front) and three csv files are saved (pose estimation, grimace scores with corresponding video frame numbers, and all video frame timestamps). The pose estimation data is directly compatible with state-of-the-art behavioural analysis pipelines and techniques such as BehaviorFlow ^33,38^.

Behind the scenes, the machine learning models perform two main tasks: pose estimation (top camera) and MGS image scoring (front camera). Pose estimation is performed using YOLO ^39^, whilst MGS image scoring is split into three tasks: identifying high quality frames suitable for MGS scoring; detecting the position of the mouse’s face; and determining the MGS score. The MGS scoring tasks are performed by three neural networks in series. The frame quality network, trained on 3200 manually annotated images and based on a modified mobilenetv3 network, scores each video frame on a continuous scale from 0 to 6 based on image quality, head position and motion blur to find the best frames for MGS scoring, The mouse face detection network, trained on 1076 manually labelled images and based on YOLOv8s ^39,40^, draws a bounding box around the mouse face, and passes the cropped image to the MGS score network. The MGS score network, trained on 1245 expert-scored images and based on a modified PyTorch Visiontransformer vit_b_16 ^41,42^ connected to two ReLU hidden layers and 5 linear activation cross-entropy output heads gives a score (0, 1, 2, or “not rateable”) for each of the five facial features (orbital tightening, nose bulge, cheek bulge, ear position and whisker change). In all results reported here, multi-fold cross validation was used to ensure that the model scoring each mouse had not been trained using any training data from that mouse, in any of the networks. For full network and training details, see methods.

### Initial validation and testing of the GrimACE (experiment 1)

In our first test of the GrimACE (experiment 1), we investigated the post-operative analgesic effects of meloxicam and buprenorphine following intracranial surgery. Mice were either unilaterally implanted with two 200μm optic fibers (one above the locus coeruleus and one within the ventral hippocampus, n=10), or were unilaterally implanted with a single 200μm optic fiber (above the locus coeruleus, n=5) (Figure 2a). To comply with the 3R principles, the choice of target sites was guided by ongoing experiments in our lab ^43^, so no animals had to be used specifically for the purpose of the experiments reported here. Approximately half of these mice received one dose of meloxicam (M) (5mg/kg s.c) shortly before surgery (n=8), and the others received 3 doses of buprenorphine + meloxicam (B+M) (0.1mg/kg + 5mg/kg s.c), once shortly before surgery, once approximately 8h after surgery, and once approximately 24h after surgery (n=7). Mice were placed in the GrimACE at various time points before and after surgery (see Figure 2b for experimental design), and 10-minute videos were recorded from front-view and top-view cameras. In addition, mice were manually inspected as part of our post-surgical cageside assessment routine, and body weight was recorded daily for 3 days following surgery.

**Figure 2.**
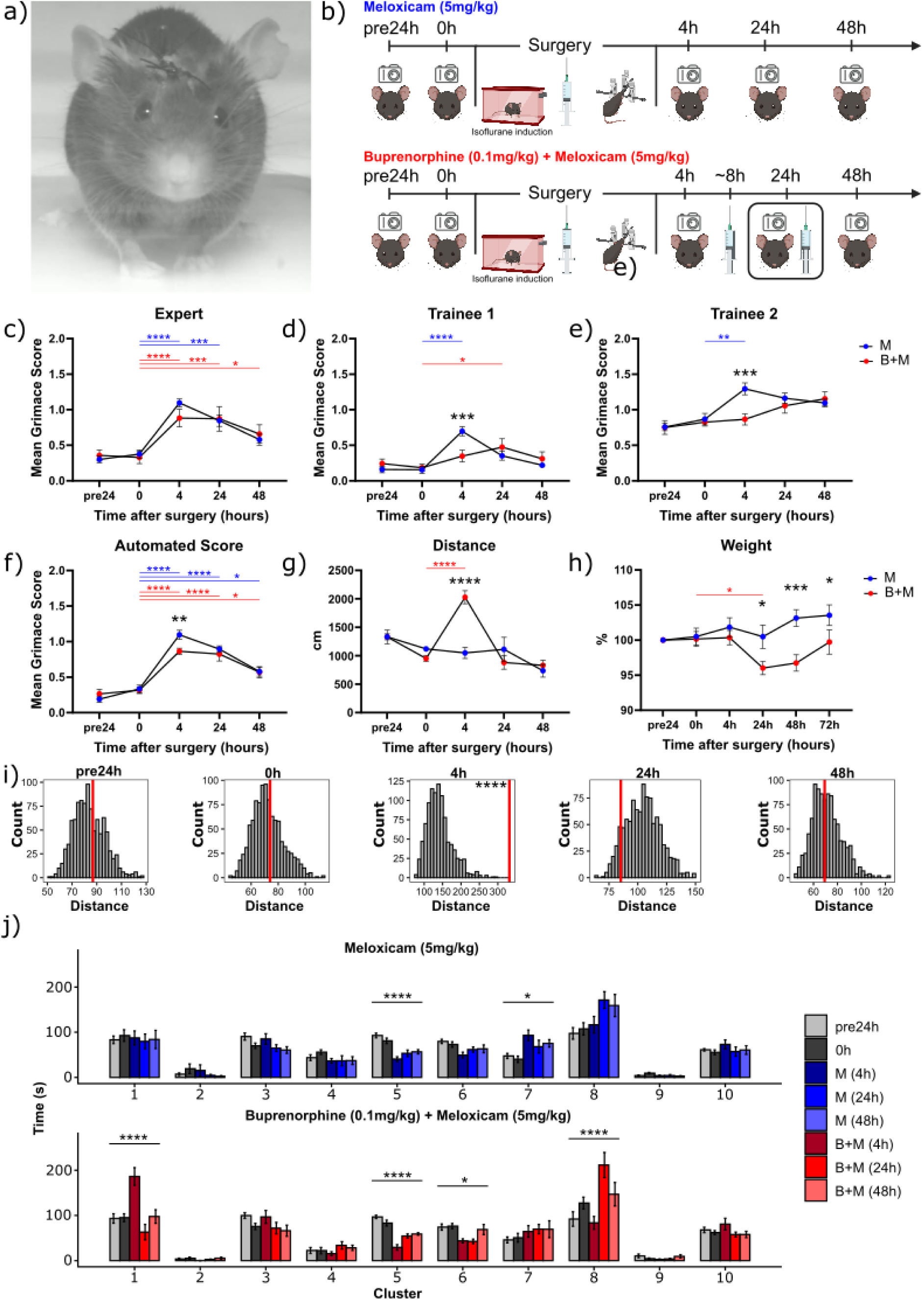
GrimACE assessment after craniotomy with two fiberoptic implants. (a) An animal 24h after surgery. (b) Experimental timeline showing GrimACE recordings, surgery and injections of analgesia (meloxicam = 8 mice, meloxicam + buprenorphine = 7 mice). (c) MGS scores from the expert rater show a significant main effect of time (F(4,50)=31.37, p=<0.0001) but no effect of treatment and no time x treatment interaction. (d) MGS scores from trainee 1 show a significant main effect of time (F(4,50)=11.41, p<0.0001), no effect of treatment, and a significant time x treatment interaction (F(4,50)=4.829, p=0.0023) (e) MGS scores from trainee 2 show significant main effects of time (F(4,63)=9.675, p<0.0001) and treatment (F(1,63)=4.551, p=0.0368), and a significant time x treatment interaction (F(4,63)=3.037, p=0.0236). (f) Automated MGS scores show a significant main effect of time (F(4,50)=64.57, p<0.0001), no effect of treatment, and no time x treatment interaction. (g) Distance moved reveals a significant main effect of time (F(4,51)=15.37, p=<0.0001), no effect treatment and a significant time x treatment interaction (F(4,51)=10.33, p=<0.0001). (h) Weight measurements show a significant main effect of time (F(5,65)=3.502, p=0.0073), no effect of treatment and a significant time x treatment interaction (F(5,65) =4.183, p=0.0023). (i) Behavior flow analysis showed no significant difference between M and B+M animals at the pre24 0h, 24h or 48h, but a significant difference between treatment groups at the 4h timepoint (p<0.0001). (n) Detailed analysis of time spent in behavioral cluster revealed a significant interaction between treatment and time in cluster 1 (2-way repeated measures ANOVA with multiple testing correction,F,(4,48)=5.236, adj p=0.03) and a significant main effect of time for clusters 5 and 7 for M animals (F,(4,24)=15.36, adj p<0.0001; F,(4,24)=5.219, adj p=0.04) and a significant main effect of time for clusters 1,5,6 and 8 in B+M animals (F,(4,24)=14.762, adj p<0.0001; F,(4,24)=31.363, adj p<0.0001; F,(4,24)=5.659, adj p=0.002; F,(4,24)=13.196, adj p<0.0001). In panels c-h, asterisks represent significant Sidak’s post hoc comparisons, smaller color-coded asterisks report drug vs time effects, larger black asterisks report between-group effects at a given time point. In panel i, asterisks represent one-tailed *z*-tests. In panel j, asterisks represent significant main effects of time in ANOVAs. *=p<0.05, **=p<0.01, ***=p<0.001, ****=p<0.0001. Error bars represent SEM.

#### Cageside Assessment

Routine cageside assessment was not able to detect signs of discomfort or pain, as all animals moved normally, their coats were well-groomed, they did not show overt postural changes, and wounds were closed and healed normally. On our cageside assessment scale, which ranges from 0 (normal) to 3 (severely affected) for several measures (see methods for details and Supplementary Figure 1 for an example of a cageside assessment score sheet), all animals consistently scored 0 on all measures. This is in line with our extensive experience performing craniotomies in the lab ^43–47^ and with the notion that craniotomy leads only to moderate pain levels that can be well managed with M or B+M. However, recent work has clearly shown that signs of pain can be detected even with adequate analgesia regimes in the hours and days following brain surgeries in mice ^27,29^. Therefore, we also investigated post-surgical pain using the MGS.

#### Mouse Grimace Scale Scoring

From the front-view video recordings collected in the GrimACE system we manually selected 10 frames from each 10-minute video for manual grimace scoring by two trainees and one expert. The expert had been scoring and providing training on the MGS for years and had also trained both of the trainees (see methods for details). Before the images of the present project were evaluated, Trainee 1 had only completed the MGS training. In contrast, in addition to the training, Trainee 2 had already scored images of C57BL/6J mice and BALB/c mice in two previous projects. Notably, we found high inter-rater variability (Supplementary Figure 2a), but very high intra-rater correlations (Supplementary Figure 2b-d). In contrast to our cageside assessment, all raters detected an increase in MGS scores of some animals at the 4h timepoint in comparison to the 0h baseline recordings. Four hours after surgery, both trainees detected significantly higher pain levels in M mice, while the expert rater did not, although the expert did rate values for M nominally higher than for B+M (Figure 2c,d,e). Interestingly all the raters still reported slightly elevated MGS scores even 48h after surgery. The automated MGS score was similarly able to resolve the surgery-induced peak in pain levels at the 4h time point (Figure 2f), identified a significant difference between M and B+M at this time point, and detected slightly elevated MGS scores in both groups at 48 hours post surgery.

#### Body weight and pose estimation

Daily body weight measurements revealed a significantly lower body weight for the B+M group compared to the M group at the 24h, 48h and 72h timepoints (Figure 2h). However this drop in body weight was in the range of 5%, which is far from our termination criterion of 85% body weight loss. Pose estimation from the top-view camera enabled simple tracking of locomotor activity (raw distance ^38^). At the 4h time point, B+M mice move significantly more than M mice (Figure 2g). This is consistent with a large body of literature showing that acute administration of opioids in general, and buprenorphine specifically, induces hyperactivity in mice and rats ^48,49^. The sharp increase in locomotor activity during the acute postoperative period when rest would be required, and the concomitant reduction in body weight, together point towards unwanted side-effects of opioid treatment that likely hamper the recovery process.

#### BehaviorFlow analysis

Top-view pose estimation also enabled more advanced analysis of behavioral motifs and the dynamic transitions between these motifs, using BehaviorFlow ^33^. BehaviorFlow was developed with the aim to recognize subtle alterations in the microstructure of mouse behavior, to reveal latent phenotypes that would otherwise remain hidden from human observers. To test whether the two analgesia regimes differentially impacted the animals, we split behavior observations into 10 clusters and compared the BehaviorFlow between both groups at every time point (see methods). Analysis indicates that at the 4h timepoint M and B+M groups showed highly significant differences in their behaviour flow (Figure 2i), which is in line with the spike in locomotor activity observed in the B+M group at that timepoint. For the remaining timepoints, no differences were detected, indicating a similar behavior flow for the two groups. We then quantified the time spent in each behaviour cluster (Figure 2j). Only cluster 1 showed a significant interaction between treatment and time. In addition, we found significant main effects of time for clusters 5 and 7 in M animals, and significant main effects of time for clusters 1,5,6 and 8 in the B+M animals. While the overall BehaviorFlow analysis (Figure 2i) corroborates the distance result (Figure 2g), we see that not every aspect of the animals’ behavior is completely dominated by the increased distance travelled by the B+M mice, which implies that detailed analysis such as those presented here could be more informative when looking for less obvious behavioural differences in other circumstances. We provide a detailed evaluation of the clusters in the discussion.

### Replicated validation of the GrimACE (experiment 2)

We were surprised to see the vast differences between human raters, and that for some raters the pain levels seemed to persist, albeit at a very moderate level, 48 hours after surgery. This contrasts with previous work using the MGS after craniotomy, which found that pain levels return to baseline after 48 hours even without analgesia (Che et al 2019). We therefore repeated the experiment to validate our findings and again investigated the post-operative analgesic effects of M and B+M following intracranial surgery, however, this time the mice underwent surgery to implant a bilateral hippocampal cannula (Figure 3a). The reason was that, in keeping with 3R guidelines, we used animals that were scheduled for ongoing experiments in our lab ^50^, thus reducing the amount of animals needed to test and optimize the GrimACE system. We also included an additional monitoring timepoint 72h post surgery (see Figure 3b for experimental design).

**Figure 3.**
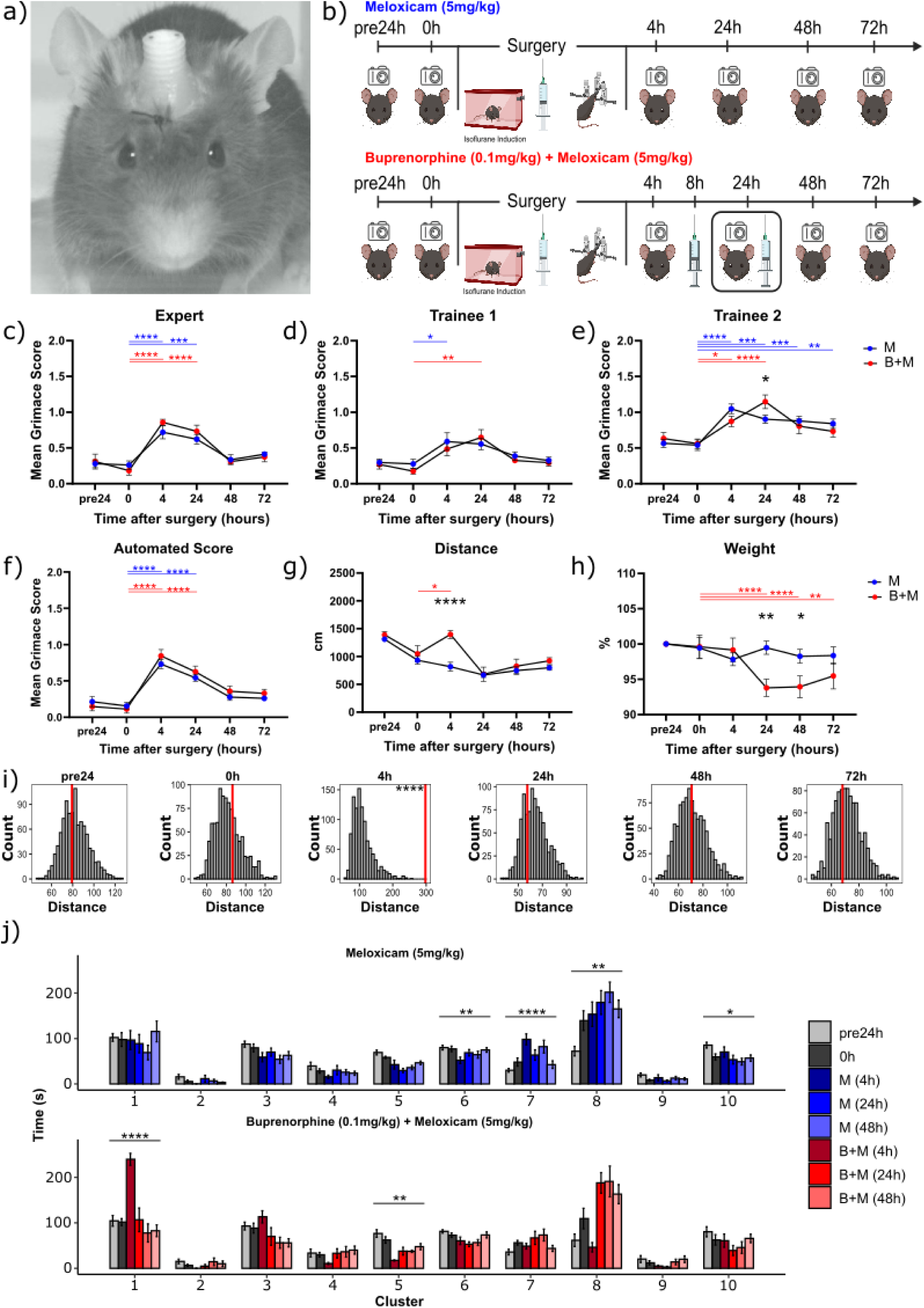
GrimACE assessment after craniotomy with cannula implantation. (a) An animal 24h after surgery. (b) Experimental timeline showing GrimACE recordings, surgery and injections of analgesia (meloxicam = 10 mice, meloxicam + buprenorphine = 6 mice). (c) MGS scores from the expert rater show a significant main effect of time (F(5,69)=24.44, p<0.0001) but no effect of treatment and no time x treatment interaction. (d) MGS scores from trainee 1 also show a significant main effect of time (F(5,69)=7.699, p<0.0001), no effect of treatment, and no time x treatment interaction (e) MGS scores from trainee 2 show significant main effects of time (F(5,69)=17.64, p<0.0001) and a significant time x treatment interaction (F(5,69)=2.681, p=0.0284). (f) Automated MGS scores show a significant main effect of time (F(5,69)=44.80, p<0.0001), no effect of treatment, and no time x treatment interaction. (g) Distance moved reveals a significant main effect of time (F(5,70)=22.12, p<0.0001), treatment (F(1,14)=7.757, p=0.0146) and a time x treatment interaction (F(5,70)=4.006, p=0.0030). (h) Weight measurements show a significant main effect of time (F(5,70)=9.813, p<0.0001), no effect of treatment and a significant time x treatment interaction (F(1,14)=1.694, p<0.0001). (i) Behavior flow analysis showed no significant difference between meloxicam and buprenorphine + meloxicam animals at the pre24, 0h, 24h, or 48h, but a significant difference between treatment groups at the 4h time point (p<0.0001). (j) detailed analysis of time spent in behavioral cluster reports a significant interaction between treatment and time only for cluster 1 (2-way repeated measures ANOVA with multiple testing correction, F(5,70)=6.86, adj p=0.000876), and a significant main effect of time in clusters 6, 7, 8 and 10 for meloxicam animals (F,(5,45)=4.75, adj p=0.01; F,(5,45)=9.993, adj p<0.0001, F,(5,45)=5.435, adj p=0.0054, F,(5,45)=9.993, adj p<0.0001) and a significant main effect of time in clusters 1 and 5 in buprenorphine and meloxicam animals (F,(5,25)=14.357, adj p<0.0001; F,(5,25)=12.41, adj p=0.00429). In panels c-h, asterisks represent significant Sidak’s post hoc comparisons, smaller color-coded asterisks report drug vs time effects, larger black asterisks report between-group effects at a given time point. In panel i, asterisks represent one-tailed *z*-tests. In panel j, asterisks represent significant main effects of time in ANOVAs. *=p<0.05, **=p<0.01, ***=p<0.001, ****=p<0.0001. Error bars represent SEM.

#### Mouse Grimace Scale Scoring

The same three raters again manually scored the extracted frames, and again all raters detected an increase in MGS scores of the animals at the 4h time point in comparison to the 0h time point (Figure 3c,d,e). Again, MGS scores remained elevated for 24 hours. This time, no differences emerged between M and B+M groups, apart from at the 24h time point according to Trainee 2. Trainee 2 also reported that the MGS scores for the M animals were raised at both the 48 and 72h timepoints, however, both the Expert and Trainee 1 report that at both 48 and 72 hours after surgery pain levels had returned to baseline. The automated MGS score also detected an increase in the MGS score at the 4h timepoint in both groups which persisted throughout the 24h timepoint, before returning to baseline at the 48 and 72h timepoints (Figure 3f).

#### Body weight and pose estimation

Similarly to experiment 1, we detect a significant difference in locomotor activity at the 4h timepoint between the M and B+M group (Figure 3g). The weight of the B+M group again dropped significantly at the 24h and 48h timepoints (Figure 3h). Again, the reduction in body weight was modest and stayed between 5-10%, before returning to baseline.

#### BehaviorFlow Analysis

BehaviorFlow analysis (Figure 3i) again showed that the M and B+M groups had significant differences in their behaviour flow only at the 4h timepoint, indicating that BehaviorFlow differences are in line with the notable spike in locomotor activity. Detailed analysis of time spent in each behaviour cluster again showed a significant interaction between treatment and time only for cluster 1, (Figure 3j). As in Experiment 1, this effect was driven by a sharp peak of time spent in cluster 1 at the 4h timepoint in the B+M group, but not in the meloxicam group. In addition, significant main effects of time emerged for clusters 6,7,8 and 10 in M animals, and for clusters 1 and 5 in B+M animals.

### No-surgery control for repeated testing (experiment 3)

The behavioral changes observed in response to buprenorphine injection were dramatic, yet the quantification of different behavior clusters also revealed considerable fluctuations across timepoints in both treatment groups. To ensure that the MGS and behavioural changes we observed in experiments 1 and 2 arose from the treatments and not from habituation, we performed a no-surgery control experiment. We repeated the exact experimental design of GrimACE exposures across 5 consecutive days as in the previous experiments (Figure 4b), but without surgery or drug administration (n=8). The results from experiment 3 show that the GrimACE reports stable and pain-free MGS values for these control animals across all timepoints (Figure 4c). However, locomotor activity slightly decreases over time (Figure 4d), similarly to what we observed in M treated animals in experiments 1+2. Body weight also remains stable (Figure 4e). Behavioral clustering revealed some fluctuations of behavior across days, but the clusters that changed significantly over time in the M and B + M treatment groups in experiment 1+2 (clusters 1,6,7,8 and 10) did not change significantly over time in this control cohort which implies that this could have been due to their treatment. Whilst these data suggest that the mice do habituate to the GrimACE, the consistent MGS scores and behaviour flow data suggest that habituation to the GrimACE setup is not necessary.

**Figure 4:**
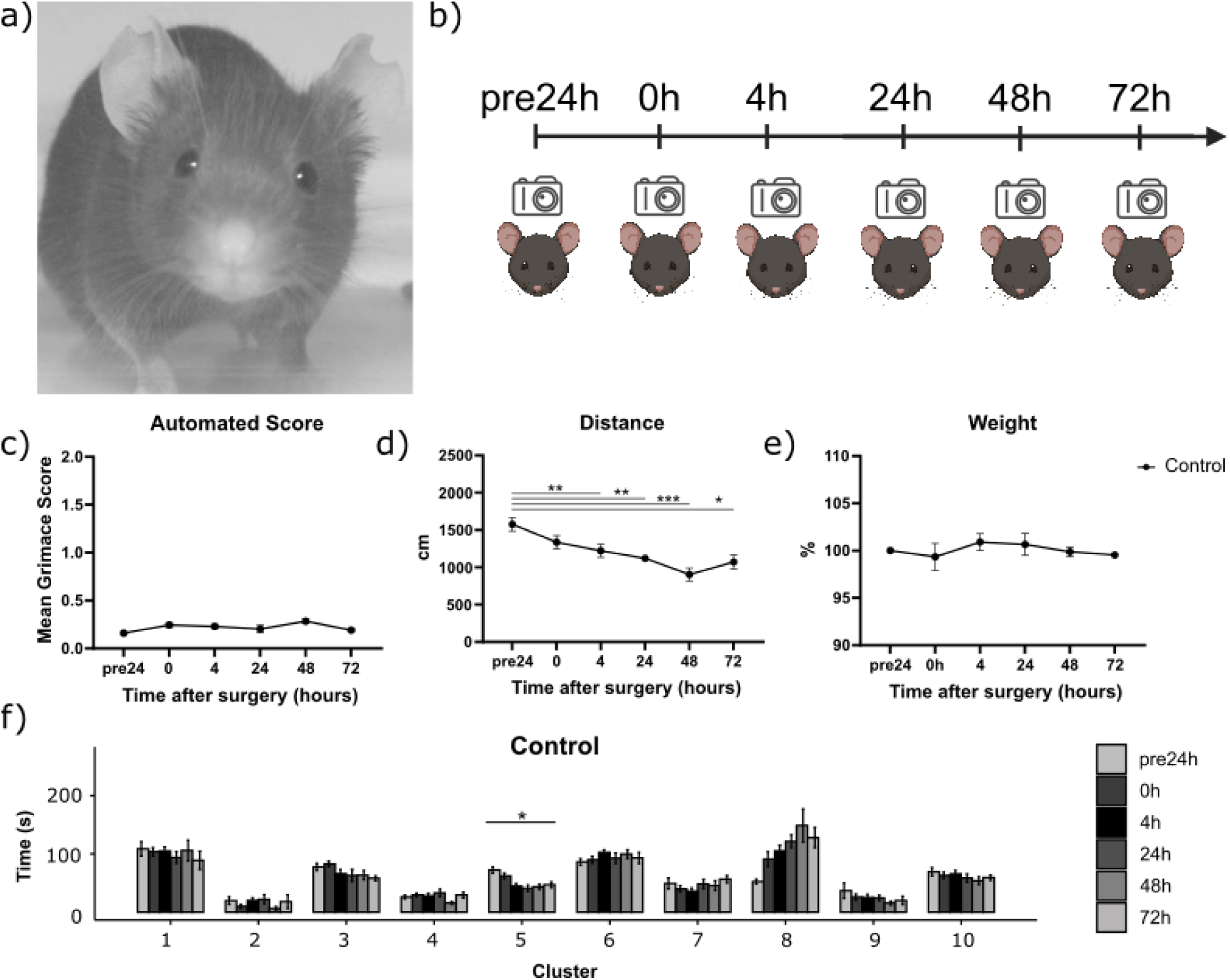
No-surgery control shows stable pain-free MGS values. (a) a control animal in the GrimACE at the 24h time point. (b) Experimental timeline showing control GrimACE recordings matched to timepoints in experiments 1 and 2. (c) no significant changes in MGS score were observed. (d) Distance reveals a significant main effect of time (F(3.145,22.01)=12.56, p<0.0001), and individual (F(7,35)=4.456, p=0.0012). (e) Weight remains stable. (f) Time-in-cluster analysis shows significant changes for cluster 5 (F(2.9,17.41=13.034, p=0.000115,) which might be indicative of arena habituation. In panel d, asterisks represent significant Tukey’s post hoc comparisons. In panel f, asterisks represent significant main effects of time in ANOVAs. *=p<0.05, **=p<0.01, ***=p<0.001, ****=p<0.0001. Error bars represent SEM.

### Automated vs Manual performance

The trainee raters gave remarkably different MGS scores from the expert and from each other on the same set of images (Supplementary Figure 2a). For this reason, only the expert rater was used to train the automated MGS scoring network, and indeed this network shows impressively consistent performance with the expert rater in leave-one-animal-out cross validation (Figure 5). When assessed on a per-video basis, the overall automated MGS score shows minimal bias across all animals (Figure 5a). Individual component scores suffer from a lack of high MGS training examples (i.e. few examples where the feature indicates severe pain and is scored as 2), which leads to an overall low-score bias of some component scores (Figure 5d-f). Whilst we chose to train the network on the data from the expert in these experiments, all of the raters showed a high degree of self-consistency (Supplementary Figure 2, b-d), which suggests that the MGS scoring network could be trained to replicate the performance of any of these raters. Fine-grained self-consistency of human raters across the individual facial components of the MGS in individual images was also consistent for features such as orbital tightening and ear position, but was more highly variable for features such as nose and cheek bulge and whiskers (Supplementary Figure 3). When comparing the individual feature scores of the expert to the automated rater in the context of experiment 1 and 2, we see that the automated rater performs similarly in the majority of features, with the largest differences being in nose and cheek bulge in experiment 2, this is likely due to the lower intra-rater correlation of these features (Supplementary Figure 4). It is also worth noting that the ear position score remains high following surgery, which could be related to the surgery altering the position of the ears. With more training data this effect could be circumvented if it is indeed related to the surgery and not pain.

**Figure 5:**
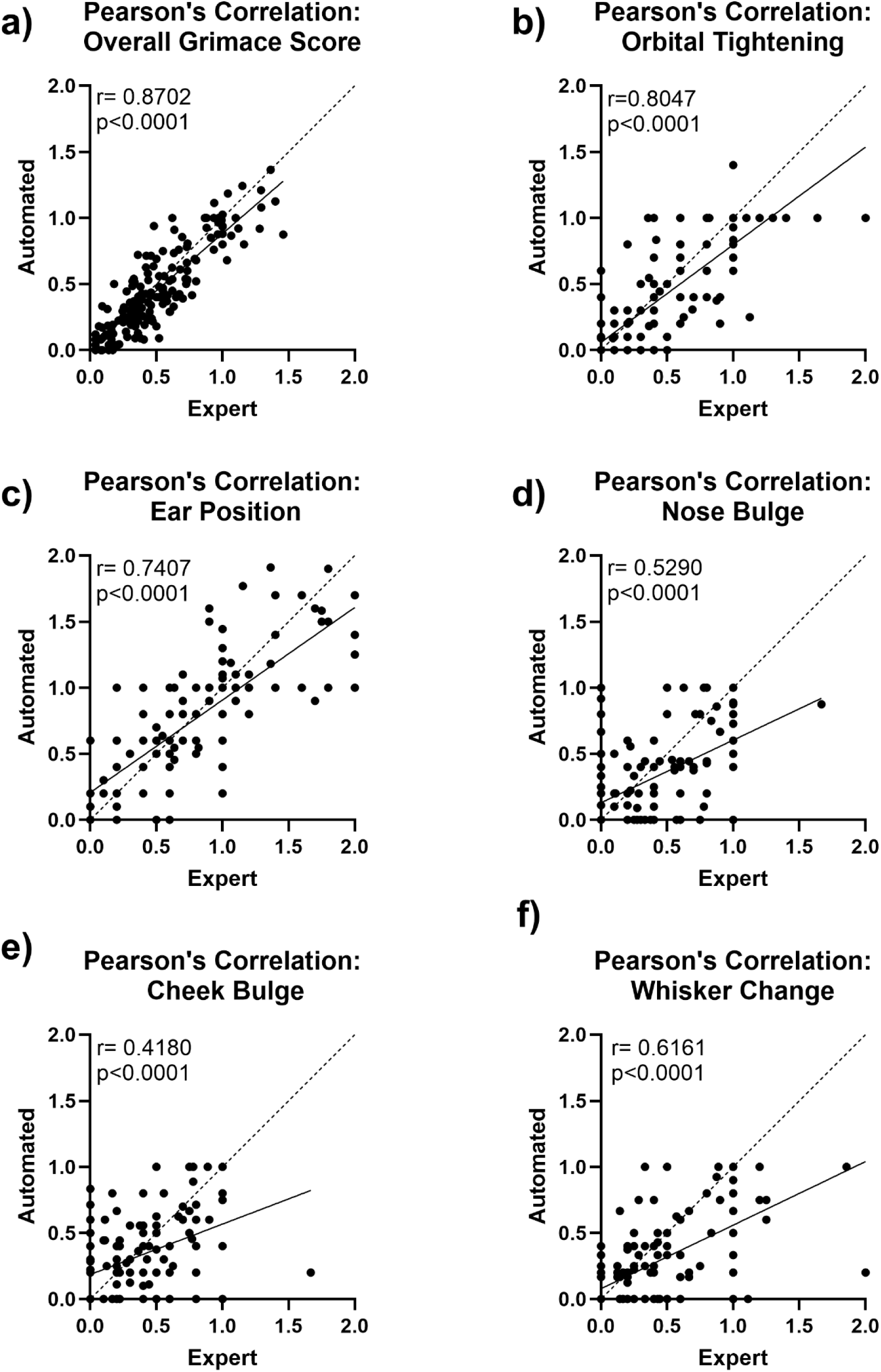
Automated MGS scoring performance. (a) Mean automated MGS scores for each feature in experiments 1 + 2 show a remarkably high overall correlation with the expert rater (Pearson’s r=0.87). Breaking these scores down into their mean components (b-f) demonstrates that the neural network is successfully distinguishing individual MGS features, where only cheek correlation has Pearson’s r < 0.5 (e). Generally, the automated MGS scores suffer most from a lack of training data with high MGS scores (severe pain levels), particularly for nose, cheek, and whisker (d-f), leading to an MGS score underestimation bias in these areas.

## DISCUSSION

### Standardization and automation of holistic welfare monitoring

Measuring spontaneous pain and discomfort in laboratory rodents is surprisingly hard ^2,19^. Although the use of facial features holds great promise for pain assessment across species, the excitement is hampered by the notion that MGS scores should not be interpreted in isolation ^2^, and the fact that MGS scoring is quite difficult and requires significant investment in training personnel to overcome high inter-rater variability ^21,51,52^. Here, we address both of these problems by combining MGS scoring with in-depth analysis of body posture and behavioral flow, and by standardizing the recording setup to train a machine learning agent on reproducibly high-quality images. We show that the combination of automated MGS scoring with simple movement tracking and advanced behavioral profiling reveals moderate increases in pain scores after craniotomies, when standard cage-side assessment fails to detect overt signs of pain or discomfort, and that it reveals known side-effect profiles of acute opioid administration after surgery. Additionally, we observe that the pain scores are slightly higher in experiment 1 (Figure 2) than experiment 2 (Figure 3). Whilst this could be due to the different implants, it could also be due to less obvious confounders. The surgeries in experiment 2 were performed by a more experienced surgeon which likely reduced the duration of the surgeries. Experiment 1 was also carried out by a female experimenter, whereas experiment 2 was carried out by two male experimenters, previous work has shown that this could be the reason for the reduced pain scores observed here ^53^.

#### Automation of MGS assessment

The automation of MGS scoring can be broken down into two parts: image acquisition automation, and image assessment automation. Previous work has focused on the automation of image assessment, and laid the foundations for the present work, by demonstrating that automatic machine learning approaches were feasible, powerful and accurate alternatives to human MGS scoring. Tuttle et al (2018) ^22^ used the (then) state of the art InceptionV3 CNN architecture to train a binary classifier targeting to distinguish “pain” from “not pain”, and showed that output confidence correlated strongly with total MGS score. Andresen et al (2020) ^25^ explored several CNN architectures, with and without pre-training, and achieved a binary classification accuracy of up to 99%. McCoy et al (2024) ^24^ took these approaches another step further by embedding CNN-based MGS scoring into a full online PainFace MGS data platform, making these automation approaches more accessible than ever, facilitating the collection of MGS training data, and demonstrating impressive performance. The fundamental difference between the GrimACE and these previous works is that it is the first end-to-end fully automated system operating in research use, encompassing image acquisition. The standardisation of image acquisition can have dramatic impacts on all subsequent steps in MGS assessment. This standardisation may be a contributing factor in the remarkable self-consistency of even trainee MGS raters, despite showing strong differences from an expert rater. This is why project-specific training of human raters may be beneficial, so that greater consensus can be achieved. Because of these fundamental improvements in training and validation data, we are reluctant to make strong claims about relative accuracy, although our overall MGS Pearson’s r of 0.87 is certainly state of the art. Unlike previous approaches, this accuracy leverages vision transformers, rather than traditional CNN architectures, which permits rapid transfer learning with small training datasets. Generally though, we emphasise that open-source computer vision tools have reached a level where their application to MGS will primarily be limited by the quantity and quality of training data, and any published network architecture can therefore be expected to improve in performance over time as new training data become available.

There are several advantages of the standardised GrimACE images, specifically from a machine learning point of view. Most of these advantages stem from the fundamental limitation that without using standardised images in specialised applications like MGS, it is often infeasible to produce training datasets that adequately sample the covariance of image acquisition setups together with mouse treatments, because training images for different treatments may often be acquired by different labs. This can introduce an inescapable hidden bias that directly impacts the primary goal of many studies: treatment comparisons. Image standardisation not only removes this problem, but as a direct consequence, it also dramatically reduces the quantity of training data required for transferable performance. These two issues taken together mean that rapid dissemination of new experimental techniques (that initially are only performed in one lab) can occur in lock-step with standardised automatic pain monitoring, greatly enhancing 3R outcomes. The trade off, of course, is compatibility with other platforms (e.g. PainFace, which requires color images).

### Pain Management after craniotomies with head-implants

Currently, there are only very few studies that carefully evaluate the appropriate analgesia regime for craniotomies. For a long time, it was thought that craniotomy-induced pain is minimal, and a large number of studies even fail to report their analgesia regime in neuroscience research papers using brain surgeries for virus injections and/or head-implants ^28^. Recently, two elegant studies have been published to directly address this issue ^27,29^. First, these studies show that craniotomies (with or without placing implants on the skull) induce significant pain levels that are clearly and reproducibly captured by MGS assessment ^27,29^. However, these pain levels are moderate (typically around or below an average MGS score of 1), and these pain levels return to baseline even without analgesia two days after craniotomy ^29^. Comparing many different analgesia regimes (including meloxicam (5mg/kg, s.c.), and buprenorphine (0.1mg/kg, s.c.) to a surgery group without analgesia, the authors show that all analgesia regimes they used provide similar and effective pain relief compared to no-analgesia. However, similar to our data, pain levels are detectable 4 to 24 hours after surgery. The study by Munk et al (2024) ^27^ compared the effectiveness of NSAID-along (carprofen, 50mg/kg via drinking water) versus multimodal analgesia (carprofen + buprenorphine depot formulation) after craniotomy with relatively large head-implants for EEG recordings. They used MGS in combination with a thorough, multimodal pain assessment to show convincingly that monotherapy with NSAIDs is as effective in managing craniotomy-related pain in mice as NSAID + opioid therapy. Importantly, the slow-release formulation of buprenorphine ensured drug availability throughout the recording period in their study, whereas in our design, buprenorphine was active at the 4h time point, when pain levels are likely maximal, but at the subsequent time points we evaluated pain before the administration of additional analgesia. Thus, together with the recent work ^27,29^, these results converge on the conclusion that neither NSAID monotherapy nor the combination of meloxicam and the powerful opioid formulations can completely eliminate the remaining, moderate levels of post-surgical pain after craniotomies. In line with previous research, we show that buprenorphine, when administered acutely, causes side effects like pronounced hyperactivity, which might be detrimental during the recovery phase when rest would be recommended. The key tenet is to strike a balance between sufficient pain management and minimal administration of analgesics, to maximize pain relief while minimizing potential side effects and complications. When taking all the recent evidence into consideration, it appears in the acute phase, the first 24 hours after surgery, NSAIDS at moderate doses (meloxicam 5mg/kg or carprofen 10mg/kg) provide pain relief to some extent, and the addition of opioids provides either no or only very little additional analgesic effect. However, these findings also suggest that there might be room to improve analgesia after craniotomy, to reduce pain levels further. Given that potent depot-formulations of buprenorphine were not able to achieve additional pain relief, it is possible that pain management cannot be improved beyond this point.

#### BehaviorFlow

A more detailed look into the clusters detected by the BehaviorFlow analysis reveals that the clusters are comprised, at least partially, from identifiable behaviors such as grooming (cluster 2), clockwise or counterclockwise rotation (clusters 3 and 10), rearing and sniffing (clusters 5, 6, 9), immobility (clusters 1 and 8), head movement (cluster 7), or a combination of all of these (cluster 4). By looking at the clusters that change over time in experiment 3 (Figure 4), we can identify clusters that are involved in baseline habituation to the GrimACE. Here we see cluster 5 which contains large amounts of exploratory sniffing and interacting with or rearing against the arena. Time spent in cluster 5 appears to be high upon first exposure to the box, then much lower when the animals are in pain, or during subsequent exposures to the arena. This suggests that exploratory behaviors decrease as the animal becomes familiar with the environment ^54,55^. Conversely, cluster 8 contains a type of immobility/inactivity that appears to increase with exposure to the GrimACE, but decreases in the B+M group at a time when mice exhibit hyperactivity. Meanwhile, clusters 1, 6, 7 and 10 appear to be related to the treatment groups. Cluster 1 is highly enriched at the 4h timepoint in B+M mice, which is strange as it appears to contain yet another type of immobility/inactivity and the B+M mice are hyperactive at this timepoint. However, careful visual inspection of this cluster in both groups suggests that the B+M mice move more in short bursts, as if they want to settle but feel compelled to move frequently, whereas M mice spend more time in a different immobility cluster (cluster 8) at this timepoint. Cluster 7 appears to be more variable following M treatment, and could indicate more head movement or head specific grooming, specifically at the 4h timepoint. Cluster 6 changes over time following both M and B+M treatment, which is no surprise as it contains a relatively high amount of rearing behaviour that is likely more difficult for the mice to perform following surgery.

### Limitations

Due to the experimental needs in our laboratory, our approach focuses on brain surgeries and on C57Bl6 mice, which represent the most commonly used surgeries and by far the most frequently used mouse strain in neuroscience labs around the world ^1,56^. The current experiments were only conducted in males, because these data were collected from ongoing experiments in the lab that happened to be conducted in males. However, previous papers that used MGS in both sexes found very similar pain scores and also a very similar response to NSAID and buprenorphine injections ^27,29^. In addition, we only tested our setup on animals that received effective analgesia regimes, thus pain levels were low across animals and we do not have sufficient training data on more severe pain levels. Therefore, our setup and analysis pipeline can be readily implemented by other laboratories, but the algorithm would have to be trained and validated further on both sexes and on experimental conditions with more severe pain levels.

Our automated MGS scoring system has been designed for maximum data exploitation and flexibility, and therefore contains no hard-coded face-region biases. This means that the automated MGS scores can, without sufficient training, also take into account relevant facial features that are not codified into the MGS, such as the presence of an implant, stitches, or wet fur. We see that through sufficient context-relevant training data, the network successfully and accurately detects the decay in post surgery MGS despite the constant presence of a head implant, but for accurate results in other contexts, additional training would be required.

It is also hoped that in the future a standardised welfare monitoring system such as this one could be incorporated into the homecage of the animals so that they can be assessed in a less intrusive manner in a more ethological environment, however given the need for advances in cageside assessment right now and the difficulty of implementing this type of holistic welfare assessment in the homecage, a system such as this one can be used to provide an immediate increase in animal welfare.

## METHODS

### Animals

To comply with the 3R principles, mice for these experiments were used based on ongoing experiments in our lab ^43,50^, in keeping with the 3R principles. The cohorts assessed in these experiments happened to be males, thus we do not have data in female animals. DBHiCre mice (2-3 months of age) (experiment 1+2) and C57BL/6 mice (2-3 months of age) (experiment 3) were bred in house. Mice were maintained in a temperature (22°C)- and humidity-controlled (52 RH) OHB facility on a 12-h reversed light–dark cycle (lights on at 08:15 am) with food (M/R Haltung Extrudat, Provimi Kliba SA, Switzerland, Cat.# 3436) and water ad libitum. Mice were housed in individually ventilated cages (SealSafe PLUS GM500, Tecniplast, Germany) in groups of 3-5 per cage and used for experiments when 2–3 months old. Cages contained wood chip bedding (LIGNOCEL SELECT, J. Rettenmaier & Söhne, Germany) nesting material (tissue paper) and a red acrylic shelter. For each experiment, mice of the same age were used in all experimental groups to rule out confounding effects of age. All tests were conducted during the animals’ active (dark) phase from 9am–6pm. All procedures were carried out in accordance with Swiss cantonal regulations for animal experimentation and were approved under license ZH067/2022 and ZH001/2021.

### Stereotaxic surgeries

For experiment 1 (Figure 2) 15 male DbhiCre mice at the age of 2–3 months were subjected to stereotactic surgery. The mice were anaesthetised with 4% isoflurane (Attane^TM^, Piramal Pharma Limited, India) in a transparent induction chamber and then placed in a stereotaxic frame with continuous anaesthesia via nosecone of 2% isoflurane. For analgesia, animals in the meloxicam group received a single subcutaneous dose of meloxicam (Metacam, Boehringer-Ingelheim, Germany) in sterile saline (5mg/kg) immediately after isoflurane induction, whereas animals in the meloxicam + buprenorphine group received 3 subcutaneous injections of meloxicam + buprenorphine (Temgesic, Temgesic solution, 0.3mg/ml, Reckitt Benckiser, Switzerland) in sterile saline (5mg/kg, 0.1mg/kg respectively), one immediately after isoflurane induction, and the other two approximately 8h and 24h after surgery. Both groups received local analgesics lidocaine (2 mg/kg) (Lidocaine 1% Streuli, Streuli pharma, Switzerland) and bupivacaine (2 mg/kg) (Bupivacain Sintetica, Sintetica, Switzerland) topically applied at the site of the wound before incision. In addition, vitamin A cream (Bausch + Lomb) was applied to the eyes to stop them drying out during the surgical procedure. After the skull was exposed, bregma (defined as the intersection of the coronal and sagittal suture) was located and the skull placement corrected for tilt and scaling. Holes were then drilled either above the right ventral hippocampus (rvHC) −3.3mm AP, 3.2mm ML from bregma (n=5), or above the rvHC and the locus coeruleus −5.4 AP, 0.9 ML from bregma (n=10). Mice were then injected with either 0.25ul ssAAV-9/2-hSyn1-mCherry-WPRE-hGHp(A) 7.5 × 10E12 vg/ml (n=5 rvHC only −3.3mm AP, 3.2mm ML, −3.8mm DV), or 0.25ul of ssAAV-5/2-hEF1α/hTLV1-dlox-ChrimsonR_tdTomato(rev)-dlox-WPRE-bGHp(A) 4.7 × 10E12 vg/ml in the LC (−5.4 AP, 0.9 ML, −3.8mm DV) and ssAAV-9/2-hSyn1-GRAB_NE2m-WPRE-hGHp(A) 2.11 × 10E12 vg/ml or ssAAV-9/2-hSyn1-EGFP-WPRE-hGHp(A) 2.9 × 10E13 vg/ml in the rvHC (−3.3mm AP, 3.2mm ML, −3.8mm DV) using a pneumatic injector (IM-11-2, Narishige) and calibrated microcapillaries (cat. no. P0549, Sigma-Aldrich). An optical fiber/optical fibers (200 µm, NA = 0.37; Neurophotometrics) were then implanted 200 μm above the injection coordinates. Optical fibers were glued to the skull using a bonding agent (Etch glue, Heraeus Kulzer) and an ultraviolet-curable dental composite (Permaplast LH Flow; M+W Dental); stitches (Supramid Polyamide pseudo monofilament, non absorbable DS19, B BRAUN) were used as required.The health of all animals was monitored over the course of 3 consecutive days post-surgery using manual cage-side assessment (see below).

For experiment 2 (Figure 3) 15 male C57BL/6 mice at the age of 2–3 months were subjected to stereotactic surgery. The mice were anaesthetised with 4% isoflurane and then placed in a stereotaxic frame with continuous anaesthesia of 2% isoflurane. For analgesia, animals in the meloxicam group received a single subcutaneous dose of meloxicam (5mg/kg) immediately after isoflurane induction, whereas animals in the meloxicam + buprenorphine group received 3 subcutaneous injections of meloxicam + buprenorphine (5mg/kg, 0.1mg/kg respectively), one immediately after isoflurane induction, and the other two approximately 8h and 24h after surgery. Both groups received local analgesics lidocaine (2 mg/kg) and bupivacaine (2 mg/kg) at the site of the wound before incision. In addition, vitamin A cream (Bausch + Lomb) was applied to the eyes to stop them drying out during the surgical procedure. After the skull was exposed, bregma (defined as the intersection of the coronal and sagittal suture) was located and the skull placement corrected for tilt and scaling. Bilateral holes were drilled above the hippocampus at −1.8 mm AP and ±1.5 mm ML from bregma, followed by the implantation of a bilateral guide cannula (62036, RWD Life Science) into the dorsal hippocampus (coordinates from bregma: −1.8 mm AP, ±1.5 mm ML, −1.5 mm DV). The health of all animals was monitored over the course of 3 consecutive days post-surgery.

### Manual Grimace Scale Scoring

To obtain the ground truth and provide us with training data that could later be used to automate the grimace scoring process we first asked 3 trained experimenters to score between 5-15 images from each video and provide their grimace scores. Orbital tightening, nose bulge, cheek bulge, ear position and whisker change were all scored either 0 (not present), 1 (moderately present) or 2 (obviously present). The raters have been labeled expert, trainee 1, and trainee 2. The expert originally trained both trainees, however trainee 2 has slightly more experience than trainee 1 (i.e., she had already scored images of C57BL/6J mice and BALB/c mice in two previous projects). For the scoring process, a spreadsheet and folders containing the images were provided to the raters. All raters were blinded to the group assignment so they did not know which animal received which analgesia protocol nor which time point of the experiment they were scoring. However, the addition of the head implants makes it obvious if the image was taken before or after surgery. In some images, the fur on the mice’s faces was wet, making scoring more difficult.

### Manual Cageside Assessment Scale

The postoperative score sheet assessments range from 0 (normal) to 3 (severely affected) for several measures (see Supplementary Figure 1 for an example cageside assessment score sheet), all animals consistently scored 0 on all measures. Experiment 1 was performed by a female experimenter, whilst experiment 2 was performed by two male experimenters. Experiment 3 was performed by both male and female experimenters.

### The GrimACE

The GrimACE (Figure 1) is a complete hardware and software solution for standardised MGS image acquisition, scoring, and keypoint-based pose estimation. It consists of an aluminium extrusion frame with custom 3D printed (PLA) components to hold lights (custom IR lighting from RS Textiles), cameras (2x Basler aca-1300-60gm, 1x Computar S-Mount H0320kp lens with S-mount adapter and 1x Computar 8 mm C-Mount M0814MP2 lens), and an acrylic arena (internal dimensions 11×7×6.5cm, with an IR permeable front/lid and matte white walls). To improve standardisation and image quality matte white acrylic shields have been attached to the frame. The cameras are connected to a laptop (minimum specifications: Intel i7 cpu, NVIDIA 4070 RTX) and powered using a Netgear GS305P-200PES POE switch using 3 GigE cables. The IR light has its own 12V power supply. Once the GrimACE has been assembled and all powered components are switched on, the user should open the GrimACE app and ensure that both cameras are connected properly. Next the user should set the working directories and ensure they know which prefixes they will add to the filenames for file identification at a later stage. The user may also edit settings such as the duration of the recording, and the frame scoring interval. The user should then slide the arena out of the setup, remove the lid of the arena and place a mouse into it. The lid should be carefully closed so as to not trap the tail of the mouse. Once the mouse is safely inside the arena it should be slid gently into place as seen in Figure 1b. The user may now click start recording. As some of the components of the GrimACE are from external parties and may be discontinued, it is advised that anyone interested in using the system should contact the 3R hub at ETH Zurich (https://ethz.ch/en/research/ethics-and-animal-welfare/animal-experimentation/3r-principles/3r-hub.html), who can help with the acquisition of parts, assembly, calibration and usage of the GrimACE.

### Pose Estimation and BehaviorFlow Analysis

Pose estimation was performed using DeepLabCut ^57^ whilst the GrimACE was being developed and this was used for the analysis above. However, the GrimACE now uses YOLO ^39^ to provide real time pose estimation. Pose estimation data from the GrimACE was then processed as described in ^33,38^. In short, 41 features were computed based on DLC tracking of both mice and grimace box, including acceleration of points, angle between two point pairs, distance between two points and to nearest border, and area spanned by multiple points. These features were then normalized and expanded over ±15 frame sequences. To perform k-means clustering, we selected 18-20 recordings from each experiment at random, resulting in 56 recordings in total. The computed feature set of each frame for these recordings was then segmented into 10 clusters. A neural network was trained on these clustering results to transfer the clustering to the remainder of the recordings. Before running the behavior flow analysis, cluster labels were smoothed within a 5 frame window, followed by computing transition numbers between clusters for each animal. These transition matrices were used for the BehaviorFlow Analysis, which assesses overall differences across all transitions. A separate BehaviorFlow Analysis was performed at each timepoint comparing cluster transitions between the M and the B+M group.

### Statistical analysis

Statistical analysis was performed using Graphpad Prism 10 and R studio. In panels c-h in Figures 2,3, a mixed effects model (REML) was used to determine fixed effects of ‘Time’, ‘Treatment’ and ‘Time x Treatment’. In Figure 4 a repeated measures one-way ANOVA was used to determine the fixed effects of time and individual When these significant effects were determined to be significant (p<0.05), Šídák’s/Tukey’s multiple comparison test was used to further investigate the effects. Asterisks represent significant Sidak’s/Tukey’s post hoc comparisons, smaller color-coded asterisks report drug vs time effects, larger black asterisks report between-group effects at a given time point. *= p<0.05, ** = p<0.01 ***=p<0.001, ****= p<0.0001. In panel i in Figures 2 and 3, asterisks represent one-tailed *z*-tests. In panel j, of Figures 2 and 3 and panel f of Figure 4 asterisks represent significant main effects of time in ANOVAs. *=p<0.05, **=p<0.01, ***=p<0.001, ****=p<0.0001. Error bars represent SEM.

### Automated MGS scoring workflow

The MGS scoring workflow operates on contiguous segments of front camera video with a user-specified interval (e.g. 10s). Each segment of video is passed to a model that consists of three neural networks. First, a frame quality detection network selects a single frame (image) from the video segment where the mouse face is clearly visible. Next, a mouse face bounding box (MFBB) network detects the mouse face and outputs a bounding box indicating its location in the image. Then, the image is cropped around the bounding box and passed to the MGS network. The MGS network is a classifier that scores each of the orbital tightening, nose bulge, cheek bulge, ear position, and whisker change as an MGS value of 0, 1, 2, or “not rateable”.

### Frame quality detection network

The frame quality detection network was trained using 3200 images from experiments 1 and 2, which were manually scored for image quality, based on the mouse face being visible, facing toward the camera, and not being blurred by motion. Each image was assigned an overall frame-quality score between 0 and 6. The model used PyTorch Image Models ^59^ mobilenetv3_large_100 with default pretrained weights feeding into a single output with sigmoid activation. The model outputs a continuous, normalized frame quality score between 0 and 1, which is then multiplied by 6 to return it to the original frame quality scale between 0 and 6. The model was trained using Adam optimization with learning rate 0.0001 and no weight decay, and the following training augmentations: horizontal flip (p=0.5); uniform random shift in brightness (+/− 15%) and contrast (+/− 5%).

### Mouse face bounding box network

1076 images from the GrimACE front camera acquired during experiments 1 and 2, as well as other unrelated experiments were manually labelled with bounding boxes around the mouse’s face. These data were split into a 10 fold cross-validation dataset without splitting images from individual animals across folds. These training and validation sets were then used to train 10 Ultralytics YOLO v8s models with default parameter and weight settings ^39,40^. In subsequent steps, networks were chosen that were not trained using the validation animal (see below).

### MGS network

The MGS network training data consists of 1245 expert-scored images from the 31 mice used in experiments 1 and 2, with a mean of 40.16 training images per mouse (st. dev. 12.24). Because the available training data were from experiments 1 and 2, MGS network training and validation differed slightly between these experiments, and experiment 3. The MGS network training and validation for experiments 1 and 2 was conducted with leave-one-animal-out cross validation across the 31 animals from those experiments. That is, the entire MGS network was retrained from scratch and validated for each of these 31 animals independently, using only the data from the other 30 animals. The MGS network for experiment 3 was trained on all of the available training data for a fixed number of epochs, based on the observed performance of the networks for experiments 1 and 2.

The training set images for each trained MGS network underwent content augmentation by a factor of 300%: the frames immediately preceding and following the labelled training frame were added to the training set with the same MGS target values (under the assumption that the MGS score varies little within 1/30th of a second). The validation set images were not augmented in this way. Each training and validation image was then run through the mouse face bounding box network, and the image, bounding box, and target MGS score were input to the MGS network training.

The MGS network used the Pytorch Torchvision vit_b_16 vision transformer network as a base ^41,42^. Vision transformers show exceptional ability to efficiently leverage pre-trained weights in image classification tasks ^41^. We selected the versatile IMAGENET1K_V1 pre-trained weights within Pytorch Torchvision for good network flexibility in subsequent fine tuning. On top of the vision transformer, two fully connected hidden layers with ReLU activation, sizes 2048 and 1024, and input dropout rates of 0.4 and 0.2 respectively, were added. The network output had 5 linear activation heads (corresponding to orbital tightening, nose bulge, cheek bulge, ear position, and whisker change) each of size 4 (corresponding to MGS of 0, 1, 2, and “not rateable”).

During training, error was scored with a composite cross entropy loss: the sum of the cross entropy loss for each of the 5 output heads. The optimization was performed using the Pytorch stochastic gradient descent module (SGD) with initial learning rates of 0.0001 for the base vision transformer, and 0.001 for the hidden and output layers. The learning rates decayed by a factor of 0.1 every 5 epochs. Weight decay of 0 was used across all weights, and momentum of 0.95 in the SGD. Training ran for 20 epochs using a batch size of 16. Network performance plateaued after 10 epochs, so the network state after 10 epochs was used for all results.

### MGS network: Image preprocessing and training-time augmentation

Because the base vision transformer was trained on Imagenet 1K colour images, all our 1280 × 1024 greyscale input images were first broadcast to 1280 × 1024 × 3 by replicating the greyscale pixel values into each of the three “colour” channels. The training dataset images then underwent stochastic preprocessing and training-time augmentation, and the validation set images only underwent deterministic basic preprocessing.

### MGS network: Validation data pre-processing

The validation data were not subjected to any random processes or augmentation. First the bounding box was expanded by 5% about the box centre, and then squared by increasing the smallest box dimension about the centre. The image was then cropped to the bounding box, with zero padding where necessary (i.e. if the bounding box crosses the edge of the image). The cropped image pixel values were normalised using fixed mean and standard deviation calculated across the entire dataset of padded, squared, cropped images. Finally, the normalised image was resized to match the input size of the vision transformer (224 × 224 × 3) using bilinear interpolation.

### MGS network: Training data preprocessing/augmentation

For each presentation of a training image during training, the following steps were performed. The mouse face bounding box was first randomly adjusted to account for fluctuations from the mouse face detection network. With probability 0.5 each boundary of the bounding box was pushed out/in by a random amount between +/− 10% of the original box size in that dimension. With probability 0.5 it was grown by 5% in each dimension about the centre. The bounding box is then made square by growing the smallest dimension about the centre to match the largest dimension. The image is then cropped to the bounding box, and zero-padded if the bounding box crosses the image edge, to avoid introducing location bias. After cropping, the image was flipped horizontally with probability 0.5. The brightness and contrast were then randomly adjusted within the range +/−15% (brightness) and +/− 5% (contrast). The image was then either: with probability 0.25 rotated by a random angle in the range −10 degrees to +10 degrees and interpolated back to the pixel grid linearly; with probability 0.25 rotated by a random angle in the range −10 degrees to +10 degrees and interpolated back to the pixel grid using a the nearest pixel value; with probability 0.5 passed through with no rotation or interpolation. The cropped image pixel values were normalised using fixed mean and standard deviation calculated across the entire dataset (same values as for validation images). Finally, the image was rescaled to match the input size of the vision transformer (224 × 224 × 3) by interpolating either bi-linearly or with nearest pixel value, each with probability 0.5.

### Frame selection

In experiment 1, frames were selected manually for scoring. In experiment 2, SLEAP ^58^ was trained to track keypoints on the nose and ears of the mouse from the front camera. Frames where any of the nose and ear keypoints had confidence < 0.5 were excluded. Next, a binary classifier was trained on 2610 frames manually scored as either suitable for MGS or not suitable for MGS. The model used PyTorch Image Models ^59^ densenet121 with default pretrained weights and a single sigmoid output node, Adam optimisation, learning rate 0.0005, weight decay of 0.001 and no data augmentation. Frames with confidence < 0.5 according to this method were excluded. Then the frames with highest confidence were selected such that: selected frames were at least 80 frames apart; at most 5 frames per video. In experiment 3, frames were selected using the continuous frame quality detection network. From each 10 second segment of video, the frame predicted to have the highest score was selected, with a minimum quality score threshold of 4.25.

## Acknowledgements

The lab of J.B. is supported by the ETH Zurich, ETH Project Grant ETH-20 19-1, SNSF Grant 310030_204372, the Basel Research Centre for Child Health (BRCCH), the Swiss 3R Competence Center, the Hochschulmedizin Zürich Flagship project STRESS, and ERA-NET NEURON - PROGRESS (SNSF: 31NE30_219119). We thank Julia Bode for maintaining the animal colony, and the staff of the EPIC for the excellent animal care and their service to our animal facility. Parts of figures were created using Biorender.com.

## Contributions

O.S. coordinated and directed the project, wrote the manuscript with input from all authors, conducted GrimACE testing and analysis, designed and built the GrimACE, provided funding.

M.S. Designed, built and developed the GrimACE App and performed data analysis.

T.L. Performed data analysis, wrote the manuscript with input from all authors.

R.Z. Manually scored grimace images, performed surgeries and conducted GrimACE testing

M.P. Performed surgeries and conducted GrimACE testing

F.K.R. Performed data analysis

J.L., R.W., A-M.M Conducted GrimACE testing

C.B. Manually scored grimace images

K.H. Manually scored grimace images

J.B. Wrote the manuscript with input from all authors, provided the facility resources and laboratory space and supervised the project.

## SUPPLEMENTARY

**Supplementary Figure 1.**
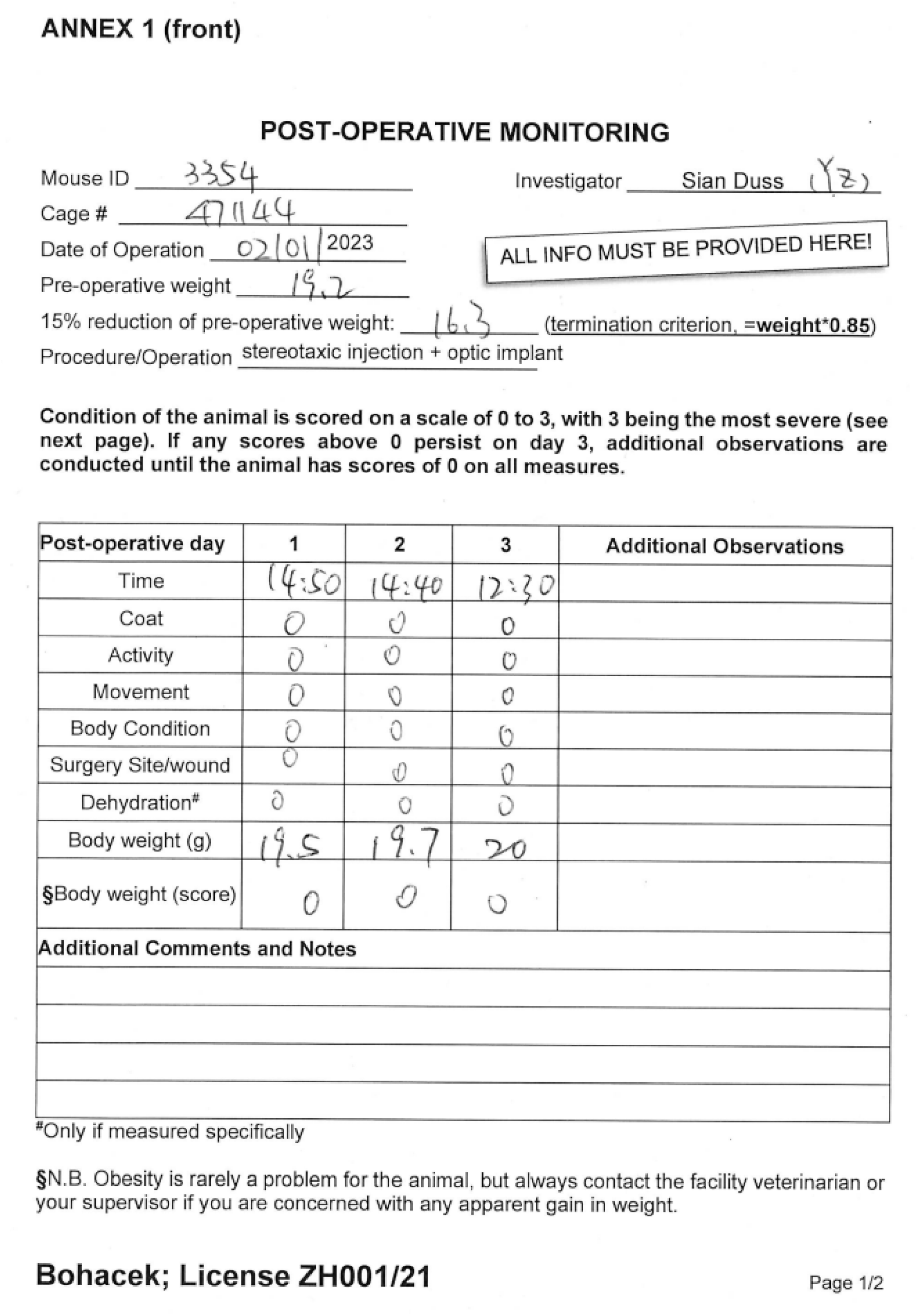
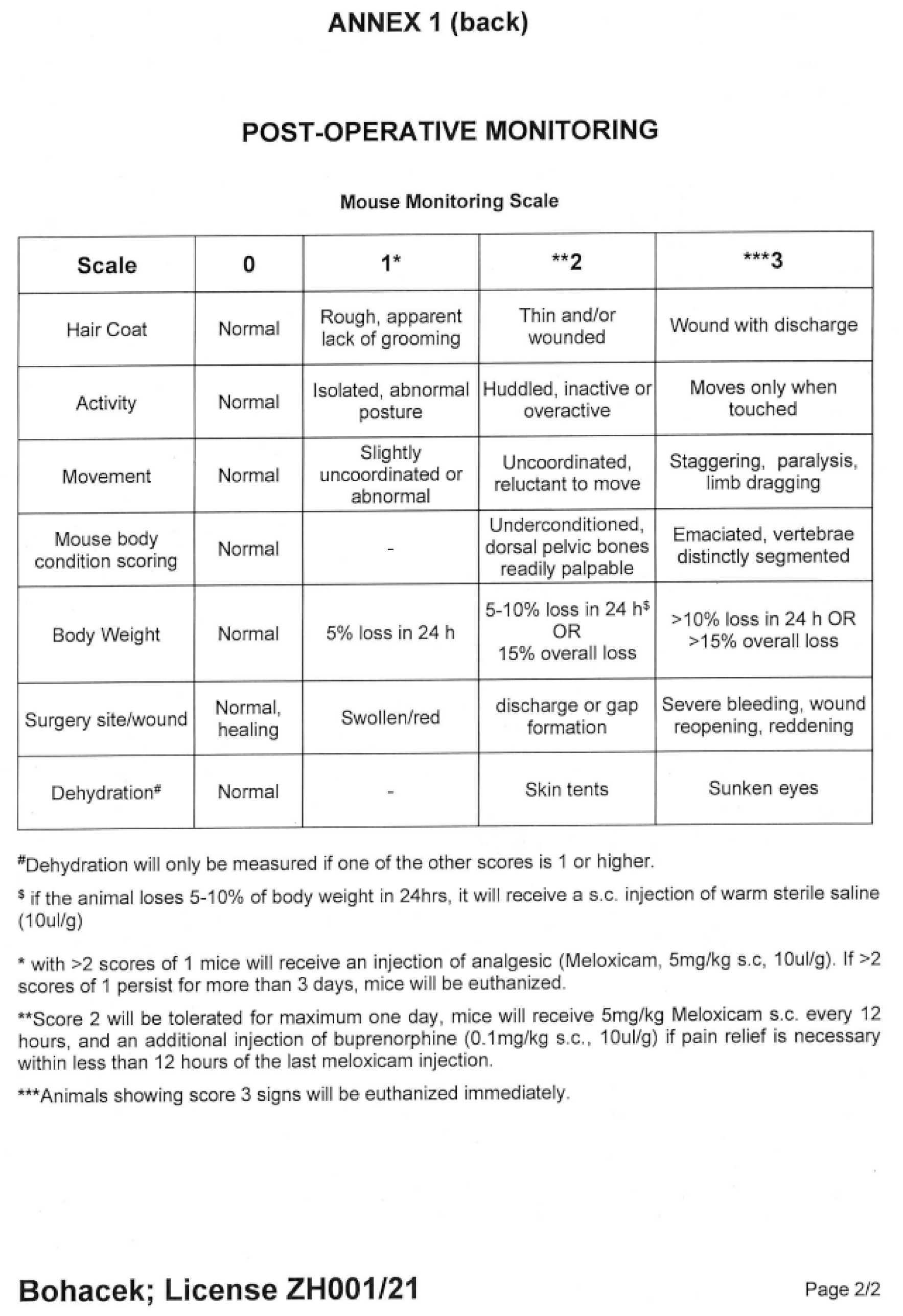
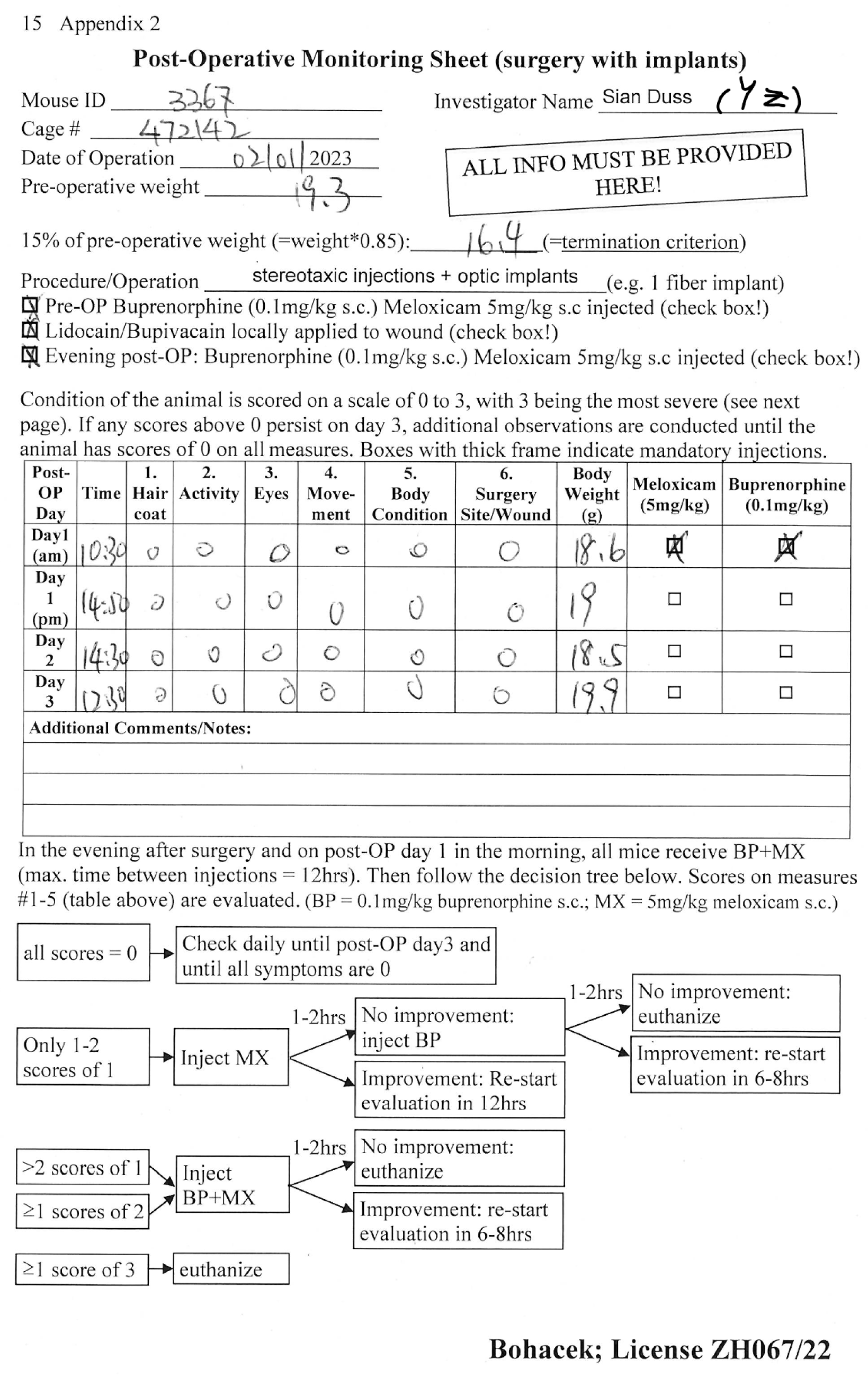
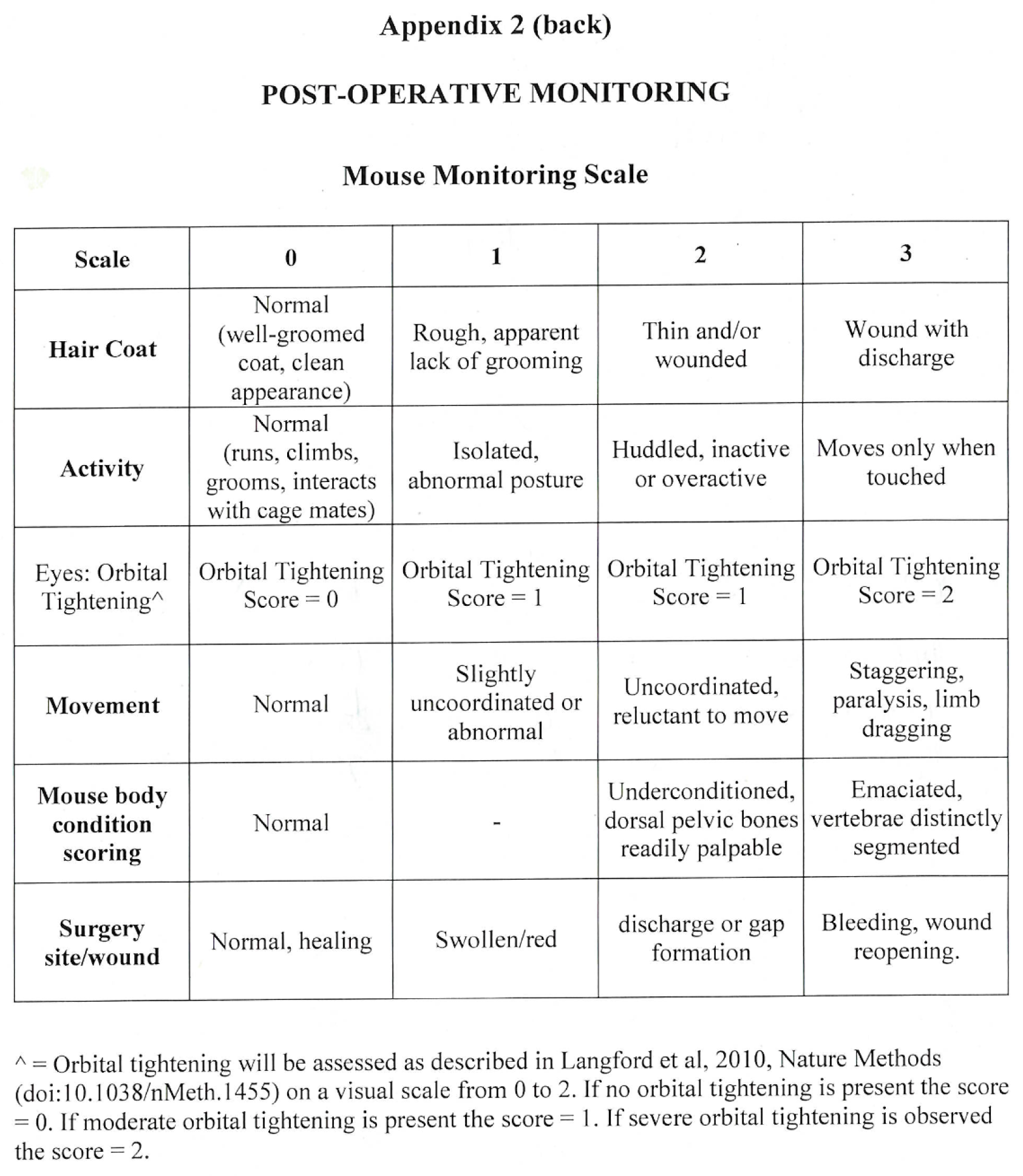
Post Operative Monitoring Scoresheets from license ZH001/21 and ZH067/22.

**Supplementary Figure 2:**
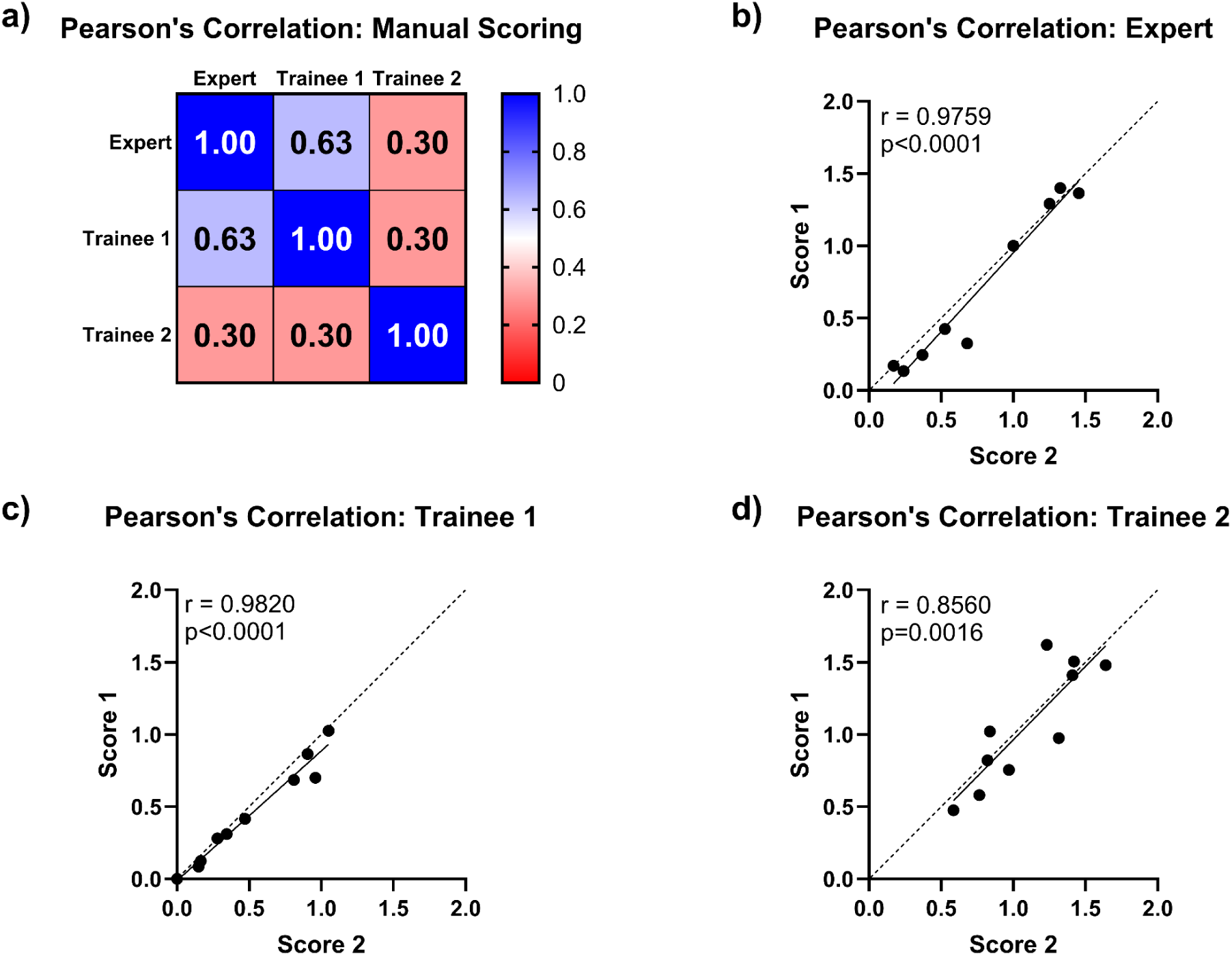
Inter-rater and intra-rater correlations for manual MGS scoring. (a) MGS scores from the three human raters show low inter-rater correlations. (b-d) Each rater scored the same 10 images twice (blinded), and the results showed highly significant intra-rater correlations, showing that each rater scored the same images with high consistency.

**Supplementary Figure 3.**
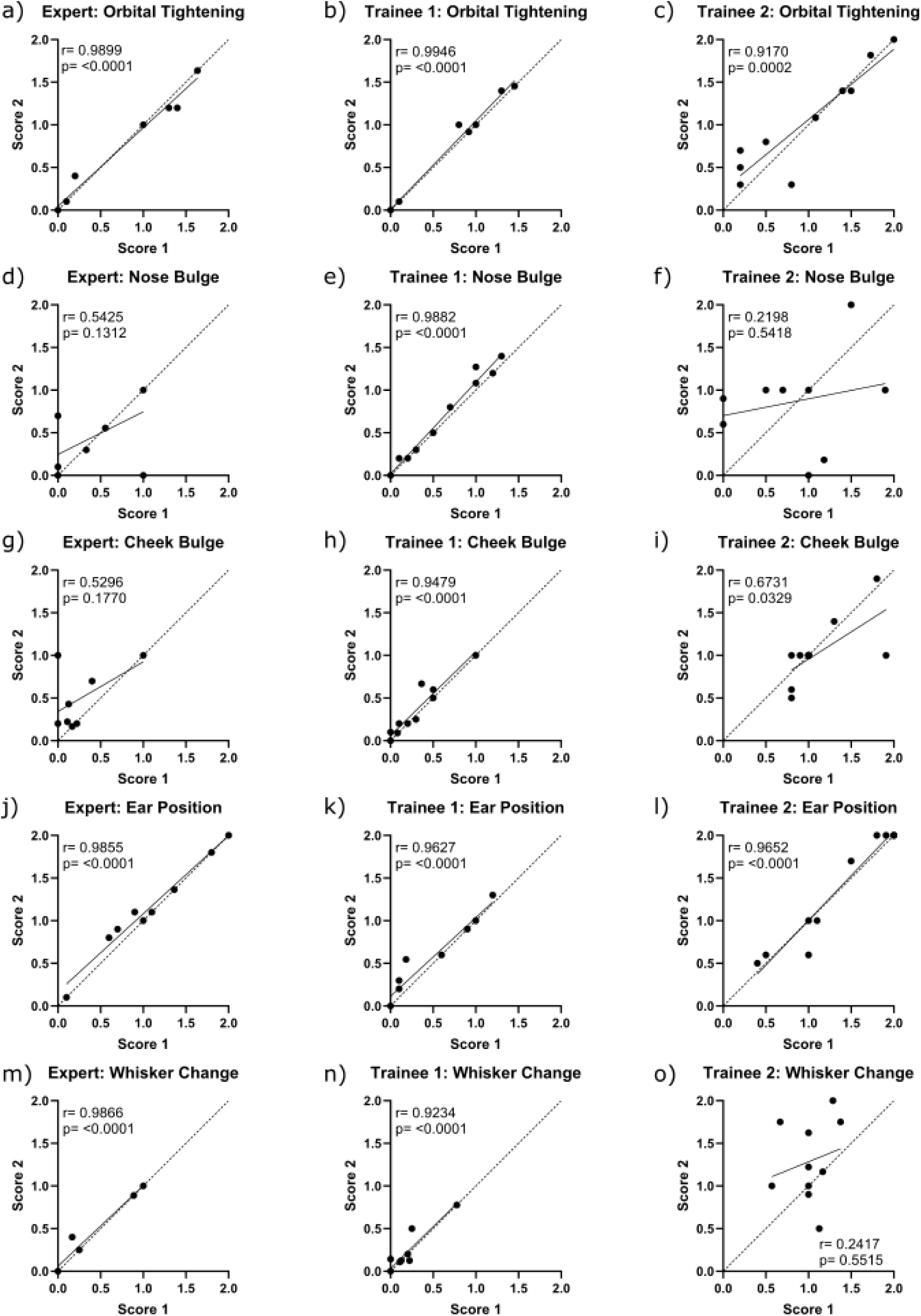
Intra-rater Pearson’s correlations for individual grimace scale features. (a-c) orbital tightening, (d-f) nose bulge, (g-i) cheek bulge, (j-l) ear position, (m-o) whisker change. Orbital tightening and ear position show high intra-rater correlations, whereas nose bulge, cheek bulge and whisker change show lower correlations for some raters.

**Supplementary Figure 4.**
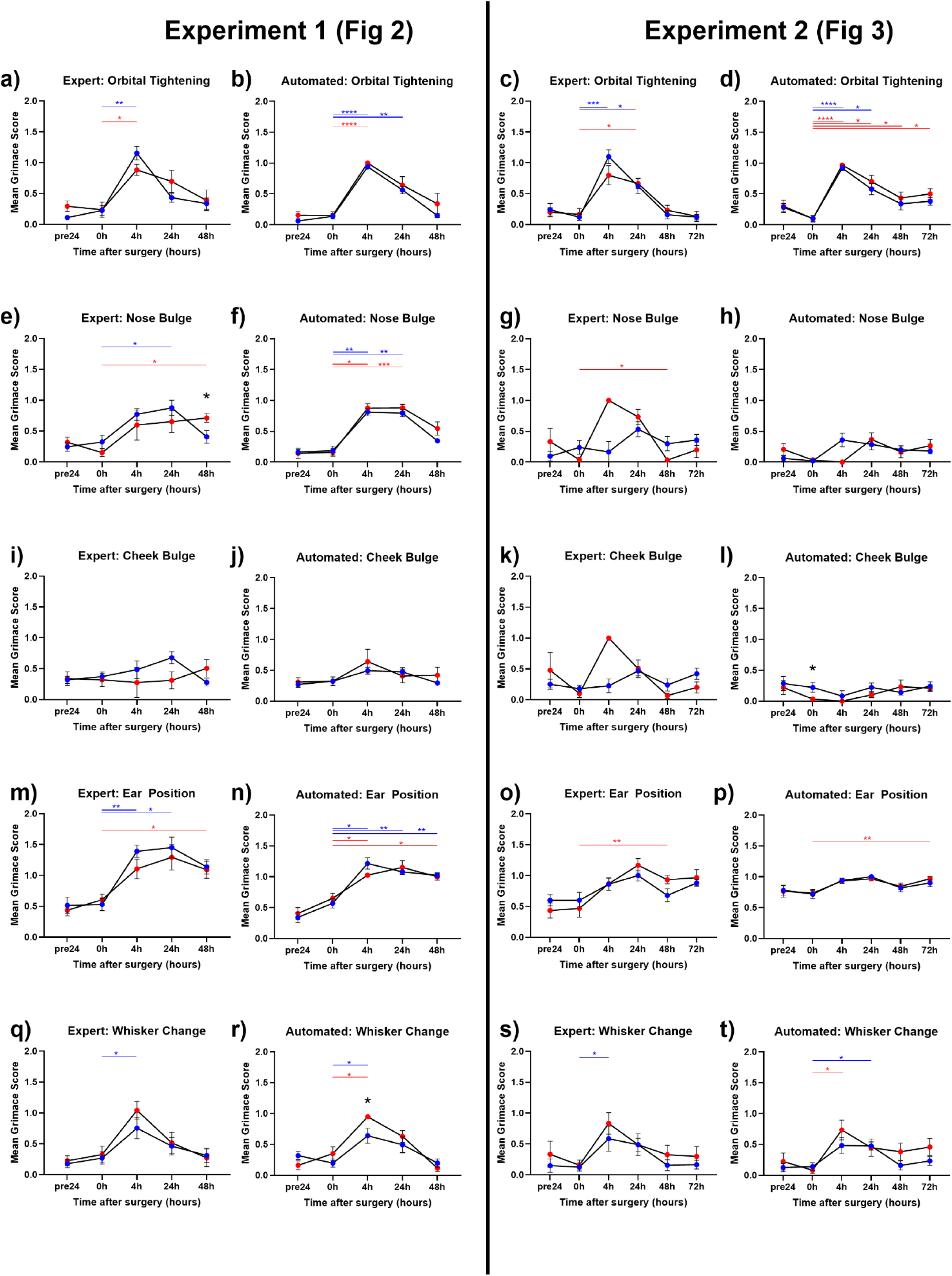
Individual grimace scale features from expert and automated raters for Figures 2 and 3. (a) Orbital tightening: MGS scores from the expert rater show a significant main effect of time (F(3.309,41.36)=29.73, p=<0.0001) and a time x treatment interaction (F(4,50)=2.666, p=0.0429), but no effect on treatment. (b) Orbital tightening: MGS scores from the automated rater show a significant main effect of time (F (2.640, 41.57) = 58.70, p=<0.0001) and treatment (F (1, 63) = 4.076, p=0.0478) but no time x treatment interaction. (c) Orbital tightening: MGS scores from the expert rater show a significant main effect of time (F(3.054,42.14)=28.39, p=<0.0001) but no effect of treatment and no time x treatment interaction.(d) Orbital tightening: MGS scores from the automated rater show a significant main effect of time (F (3.543, 48.90) = 30.90, p=<0.0001) but no effect of treatment and no time x treatment interaction. (e) Nose Bulge: MGS scores from the expert rater show a significant main effect of time (F(2.212,26.54)=9.189, p=0.0007) but no effect of treatment and no time x treatment interaction. (f) Nose Bulge: MGS scores from the automated rater show a significant main effect of time (F(3.616,55.15)=52.65, p<0.0001) but no effect of treatment and no time x treatment interaction.(g) Nose Bulge: MGS scores from the expert rater show a significant main effect of time (F(3.293,39.52)=5.685, p=0.0019) and a time x treatment interaction (F(5,60)=3.189), p=0.0128, but no effect on treatment. (h) Nose Bulge: MGS scores from the automated rater show a significant main effect of time (F(3.435,41.22)=4.565, p=0.0055) but no effect of treatment and no time x treatment interaction. (i) Cheek Bulge: MGS scores from the expert rater show no significant main effects of time or treatment, and no time x treatment interaction. (j) Cheek Bulge: MGS scores from the automated rater show a significant main effect of time (F(3.504,51.68)=3.252, p=0.0234) but no effect of treatment, and no time x treatment interaction.(k) Cheek Bulge: MGS scores from the expert rater show a significant main effect of time (F(3.201,38.41)=4.084, p=0.0116) and a time x treatment interaction (F(5,60)=2.749, p=0.0266), but no effect on treatment. (l) Cheek Bulge: MGS scores from the automated rater show no significant main effects of time or treatment, and no time x treatment interaction. (m) Ear Position: MGS scores from the expert rater show a significant main effect of time (F(2.613,32.66)=19.21, p<0.0001) but no effect of treatment and no time x treatment interaction. (n) Ear Position: MGS scores from the automated rater show a significant main effect of time (F(2.989,37.36)=44.49, p<0.0001) but no effect of treatment and no time x treatment interaction. (o) Ear Position: MGS scores from the expert rater show a significant main effect of time (F(3.817,52.67)=10.31, p<0.0001) but no effect of treatment and no time x treatment interaction. (p) Ear Position: MGS scores from the automated rater show a significant main effect of time (F(2.314,31.94)=5.558, p=0.0063) but no effect of treatment and no time x treatment interaction. (q) Whisker Change: MGS scores from the expert rater show a significant main effect of time (F(3.308,39.70)=9.685, p<0.0001) but no effect of treatment and no time x treatment interaction. (r) Whisker Change: MGS scores from the automated rater show a significant main effect of time (F(2.805,33.66)=17.64, p<0.0001) but no effect of treatment and no time x treatment interaction. (s) Whisker Change: MGS scores from the expert rater show a significant main effect of time (F(2.481,40.69)=4.760, p=0.0093) but no effect of treatment and no time x treatment interaction. (t) Whisker Change: MGS scores from the automated rater show a significant main effect of time (F(2.955,40.19)=6.810, p=0.0009) but no effect of treatment and no time x treatment interaction. In panels a-t, asterisks represent significant Tukey’s posthoc comparisons, smaller color-coded asterisks report drug vs time effects, larger black asterisks report between-group effects at a given timepoint.*=p<0.05, **=p<0.01, ***=p<0.001, ****=p<0.0001. Error bars represent SEM.

## Notes

### Competing Interest Statement

The authors have declared no competing interest.

